# Recent fluctuations in Mexican American genomes have altered the genetic architecture of biomedical traits

**DOI:** 10.1101/2020.01.13.905141

**Authors:** Melissa L. Spear, Alex Diaz-Papkovich, Elad Ziv, Joseph M. Yracheta, Simon Gravel, Dara G. Torgerson, Ryan D. Hernandez

## Abstract

People in the Americas represent a diverse group of populations with varying degrees of admixture among African, European, and Amerindigenous ancestries. In the United States, many populations with non-European ancestry remain understudied, and thus little is known about the genetic architecture of phenotypic variation in these populations. Using genome-wide genotype data from the Hispanic Community Health Study/Study of Latinos, we find that Amerindigenous ancestry has increased over time across Hispanic/Latino populations, particularly in Mexican Americans where Amerindigenous ancestry increased by an average of ∼20% over the 50-year period spanning 1940s-1990s. We find similar patterns across American cities, and replicate our observations in an independent sample of Mexican Americans. These dynamic ancestry patterns are a result of a complex interaction of several population and cultural factors, including strong ancestry-related assortative mating and subtle shifts in migration with differences in subcontinental Amerindigenous ancestry over time. These factors have shaped patterns of genetic variation, including an increase in runs of homozygosity in Amerindigenous ancestral tracts, and also influenced the genetic architecture of complex traits within the Mexican American population. We show for height, a trait correlated with ancestry, polygenic risk scores based on summary statistics from a European-based genome-wide association study perform poorly in Mexican Americans. Our findings reveal temporal changes in population structure within Hispanics/Latinos that may influence biomedical traits, demonstrating a crucial need to improve our understanding of the genetic diversity of admixed populations.

## Introduction

The United States Census Bureau refers to the Hispanic/Latino ethnicity as a self-identified category for individuals with ancestry deriving from Spain and the Spanish-speaking countries of the Americas. As such, this broad ethnic group living in the United States is a culturally, phenotypically, and genetically diverse continuum of populations. Individuals who identify as Hispanic/Latino have varying proportions of Amerindigenous, African, and European genetic ancestry, each with its own unique continental demographic history. Demographic forces such as population bottlenecks and expansions, migration and adaptation to novel environments resulted in observable differences in continental patterns of genetic variation (1–3). These differing patterns were shaped by many historical events of migration which partially included the founding of the Americas by Amerindigenous populations, the colonization by Europeans, and the African slave trade (4–8), however additional complexities surrounding these events remain highly understudied. These large-scale migrations and additional demographic events shaped the genetic diversity of individuals currently living within the United States (9–13).

Demographic history has shaped the genetic architecture of modern human phenotypic variation (14–19), and is critical to consider in the search for the genetic basis of complex diseases. The demography of the United States has changed drastically over the 20^th^ century, and by 2044 is predicted to become a ‘minority-majority’ country whereby no one racial/ethnic group comprises more than 50% of the population (20). By 2060 Hispanics/Latinos are projected to make up the largest of that share at 29% or 119 million individuals (20). However, to date, population-based medical genomics research [and its subsequent benefits, including polygenic risk score (PRS) profiling] have been disproportionately focused on individuals of European descent, with the findings primarily benefiting European populations (21, 22). Despite the increases in sample sizes, rates of discovery, and traits studied, Hispanic or Latin American participation in genome-wide association studies (GWAS) has continued to hover around 1% (23, 24). This trend, along with factors ranging from research abuse and community mistrust to community superstition and apathy have led to a situation where these populations (and other non-European populations) are particularly vulnerable to falling behind in receiving the benefits of the precision medicine revolution (22, 23).

In this study we utilize the largest genetic study of Hispanics/Latinos in the U.S. to date -- the Hispanic Community Health Study/Study of Latinos (HCHS/SOL) (10) -- to understand how patterns of genetic variation in Hispanic/Latino populations in the United States have changed over the last century, and evaluate the impact such changes may be having on complex traits.

## Results

### Global ancestry proportions among HCHS/SOL Hispanic/Latino Populations

Using the subset of sites that overlapped with our African, European, and Amerindigenous reference panels, we called 3-way global ancestry estimates for 10,268 unrelated HCHS/SOL individuals (see methods). Figure 1A summarizes the global ancestry proportions shaded by admixture estimates in a ternary plot, recapitulating the original HCHS/SOL analysis of continental ancestry (10). However, while several population groups appear to have overlapping ancestry proportions, this analysis masks more subtle structure in subcontinental ancestry. To investigate subtle population structure across these self-identified population groups, we performed UMAP on the top 3 principal components (see methods), and find substantial structure across self-identified groups (Figure 1B and Supplementary Figure 1B). We find that Dominicans, who have the highest average proportions of African ancestry, are in the middle, with Puerto Ricans and Cubans, diverging in opposite directions (Supplementary Figure 1B) with clines of increasing European ancestry proportions (Figure 1B). Further, while self-identified Mexican, Central, and South American groups appear to have overlapping ancestry proportions in Figure 1A, UMAP represents the Mexican Americans and Central/South American groups as large, separate wings that diverge from self-identified Cubans and Dominicans, with both clusters diverging with clines of increasing ancestry toward different Amerindigenous populations (Figure 1B and Supplementary Figure 1B and 1C). While UMAP places each of the Amerindigenous populations and the CEU population at the border of one or more HCHS/SOL clusters, UMAP isolates the YRI samples into a distinct island suggesting that the use of this single African population may be suboptimal for studying African ancestry in the populations sampled by HCHS/SOL (Supplementary Figure 1C).

**Figure 1.**
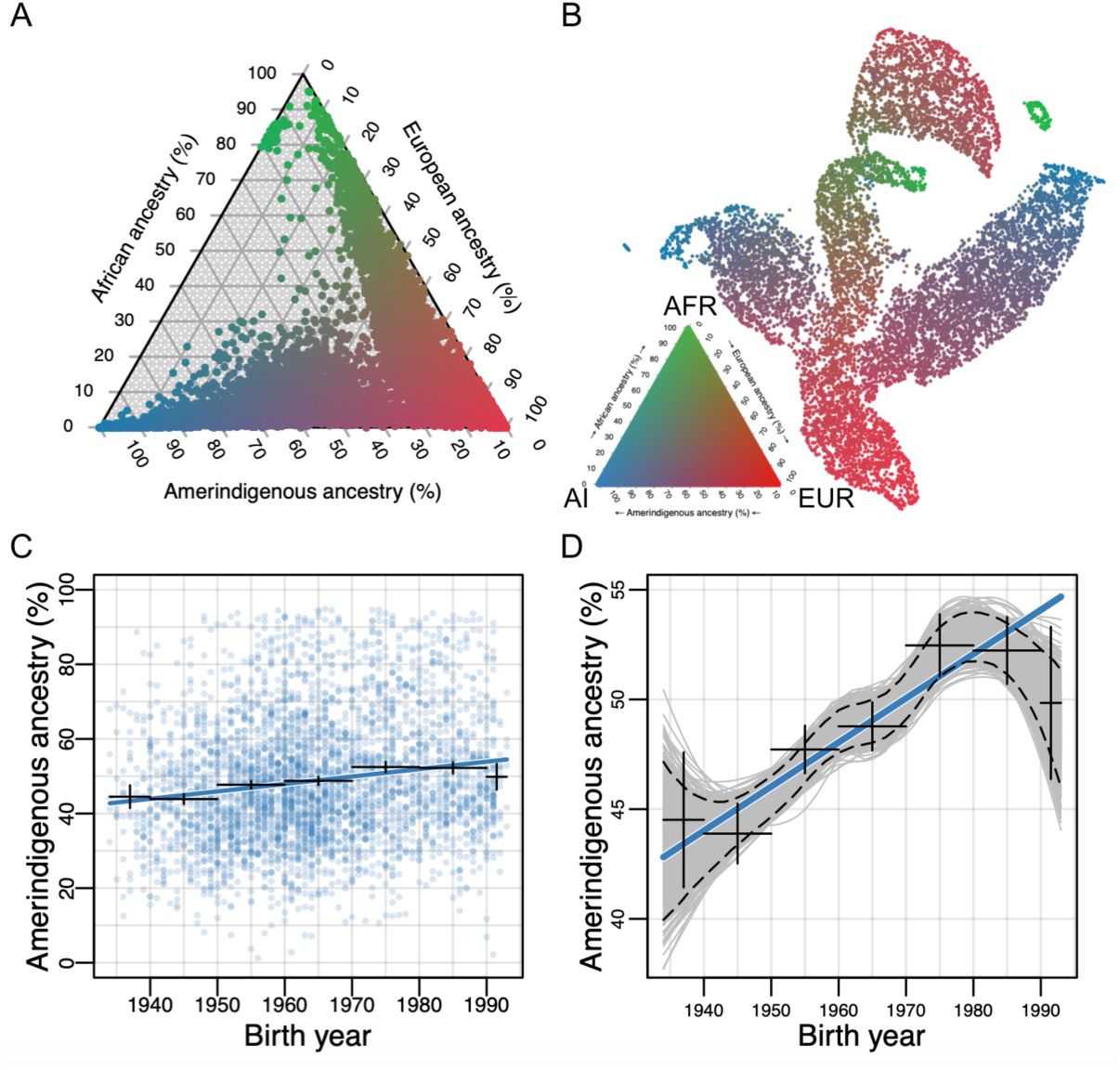
Recent dynamics continually shape the continuum of continental ancestry Hispanic/Latino populations. A. Ternary plot of HCHS/SOL (n=10,268) colored by admixture proportions. B. Uniform Manifold Approximation and Projection (UMAP) plot depicting the genetic diversity of HCHS/SOL and the reference panel (n=10,591) using 3 principal components, colored by admixture proportions (see Supplemental Fig 1 for population labels). Within the legend, AFR, EUR, and AI refer to African, European, and Amerindigenous global ancestries, respectively. C. Global Amerindigenous ancestry proportions plotted by birth year for Mexican Americans (n=3,622). Fitted line is multiple regression of Amerindigenous ∼ birth year + sampling weight. Bars represent 95% confidence intervals for individuals grouped by decade. D. Bootstrap resampling (n=1000 iterations) of Amerindigenous global ancestry for the Mexican American individuals with a fitted LOESS regression line for each iteration. Dashed lines represent the 95% confidence interval and the blue line represents the fitted regression line from Figure 1C.

### Dynamic Global Ancestry Proportions in Mexican Americans

For each of the HCHS/SOL populations, we evaluated differences in global ancestry estimates over time while accounting for the sampling method (referred to as “sampling weight”, see methods) used for the design of the HCHS/SOL study (25). We found that in all populations, the effect size for Amerindigenous ancestry on birth year is positive, though only statistically significant after multiple testing in the Mexican American (*β* =0.0023; P=3.58E-22; Figure 1C) and Central American (*β*=0.0013; P=0.0013) cohorts (Supplementary Table 1). Due to the larger sample size, magnitude of the effect, and statistical significance, we shift our focus to Mexican Americans. In Mexican Americans, the increase in Amerindigenous global ancestry over time was consistent across multiple data stratifications including recruitment region, US born or not US born, educational attainment, and sex (Table 1), and was robust to alternative methods for estimating global ancestry proportions (e.g. based on the summation of RFMix local ancestry estimates; Supplementary Figures 2 and 3). We performed bootstrap resampling (n=1000) of global Amerindigenous ancestry for the Mexican Americans and observed a consistent increase in Amerindigenous ancestry with fitted LOESS smoothing (Figure 1D) and when individuals were binned by birth year decades (Supplementary Figure 4). On average, global Amerindigenous ancestry has increased ∼20% over the last 50 years in Mexican Americans.

**Table 1:** Relationship of Amerindigenous global ancestry and birth year for Mexican Americans stratified by recruitment region, US born vs non-US born status, sex and educational attainment. For recruitment region, data stratification was limited to Chicago and San Diego as sample size for the Bronx and Miami was limited: 124 and 25 individuals, respectively. Education attainment was categorized as either less than a high school diploma or equivalent degree (<HS), equal to a high school diploma or equivalent degree (=HS), or post-secondary education (>HS).

| Category | N | Mean | Median | R2 | Effect | Std.Err | P |
| --- | --- | --- | --- | --- | --- | --- | --- |
| All | 3622 | 0.489 | 0.468 | 0.027 | 0.0023 | 0.0002 | 3.58E-22 |
| Chicago | 1310 | 0.562 | 0.550 | 0.017 | 0.0016 | 0.0005 | 0.0006 |
| San Diego | 2163 | 0.428 | 0.422 | 0.012 | 0.0012 | 0.0002 | 4.29E-07 |
| US born | 634 | 0.427 | 0.418 | 0.063 | 0.0027 | 0.0004 | 1.77E-10 |
| Not US born | 2987 | 0.502 | 0.481 | 0.050 | 0.0032 | 0.0003 | 1.38E-30 |
| Male | 1500 | 0.494 | 0.475 | 0.038 | 0.0028 | 0.0004 | 3.83E-14 |
| Female | 2122 | 0.485 | 0.462 | 0.022 | 0.0019 | 0.0003 | 3.07E-10 |
| <HS | 1518 | 0.520 | 0.500 | 0.045 | 0.0026 | 0.0004 | 1.39E-12 |
| =HS | 960 | 0.501 | 0.479 | 0.022 | 0.0018 | 0.0005 | 0.0003 |
| >HS | 1140 | 0.436 | 0.422 | 0.045 | 0.0027 | 0.0004 | 6.53E-13 |

We replicated the increase in global Amerindigenous ancestry over time in a smaller, independent cohort of self-identified Mexican Americans (n=705) from the Health and Retirement Study (HRS) [34]. The HRS Mexican Americans in this study are older compared to the HCHS/SOL Mexican Americans (birth year distribution: 1915-1981; mean=1943,median:1942) and have lower levels of global Amerindigenous ancestry on average (mean=0.29), but we still observed an increase in global Amerindigenous ancestry over time (*β*=0.00082; P=0.02; SE=0.0003673; Supplementary Figure 5A). We performed 1000 bootstrap resampling iterations of the linear regression model (global Amerindigenous ancestry ∼ birth year) fitted to the data. From these resampling iterations, 98.2% of the tests had a slope > 0 (average *β*=0.00083) and 61.5% of the regression p-values were less than 0.05 as illustrated in Supplementary Figures 5B-5D.

A previous study (12) identified ancestry biased migration in African Americans where individuals with higher proportions of European ancestry migrated first out of the South during the Great Migration followed by individuals with higher proportions African ancestry. We hypothesized that earlier immigrants to the US had higher proportions of European ancestry followed by recent immigrants having higher proportions of global Amerindigenous ancestry. In our non-US born individuals (N=2987), we evaluated differences in ancestry estimates over time while accounting for years in the US and sampling weight and identified a significant effect of years in the US (*β*=− 0.0009; P=0.0006; SE=0.0003). However, this did not change the effect of birth year on the proportion of global Amerindigenous ancestry (*β* =0.0028; P<2e-16, SE=0.0003).

For US born individuals we assessed whether parental birth place could explain the increases in global Amerindigenous ancestry. Of the 634 US born individuals, 385 had parents both born outside of the US, 149 had one parent born outside of the US. 97 had both parents born within the US. For the 385 individuals with both parents born abroad, we identified a significant association between birth year and global Amerindigenous ancestry (*β*=0.004, P=2.34e-10, SE=0.0006). For the remaining individuals we were unable to identify a significant effect of birth year on global Amerindigenous ancestry, possibly due to a small sample size.

### Individual loci are not driving global ancestry proportions

We used local ancestry estimates generated across the genome to perform admixture mapping in HCHS/SOL Mexican Americans to determine if younger individuals harbored excess Amerindigenous ancestry in certain regions of the genome. Although we tried two different models (see methods), we did not find any loci to be significantly associated with birth year across the genome (Supplementary Figure 6).

### Little evidence for subcontinental population structure

It is possible that the increase in global Amerindigenous ancestry over time could be biased by changes in the specific subcontinental Amerindigenous ancestries over time (though such an effect is not visible in our UMAP analysis, Figure 1B). If it were the case, then we would expect subtle signals of genetic divergence in Amerindigenous ancestry tracts over time. To investigate this, we calculated F_ST_ between all pairs of birth-decades (see methods). Figure 2A shows all pairwise comparisons among birth-decades, and demonstrates that while the estimates of F_ST_ are negligible (with many estimates below 0), there is a subtle trend of increasing F_ST_ as birth-decade differences increase (though individuals born in the 80s and 90s show a conflicting pattern). We further investigated this pattern using genetic diversity, π, in Amerindigenous ancestry tracts for each birth-decade (see methods). We hypothesized that if there were increased migration from multiple Amerindigenous source populations (coupled with rapid population growth in Mexican American communities), then genetic diversity should be increasing over time. We found the opposite: Supplementary Figure 7 shows a subtle decrease in genetic diversity (π) over time from the 1930s to the 1980s in non-US born Mexican Americans, and a subtle decrease in US born Mexican Americans from the 70s to the 90s (while remaining roughly constant from the 30s to the 70s).

**Figure 2.**
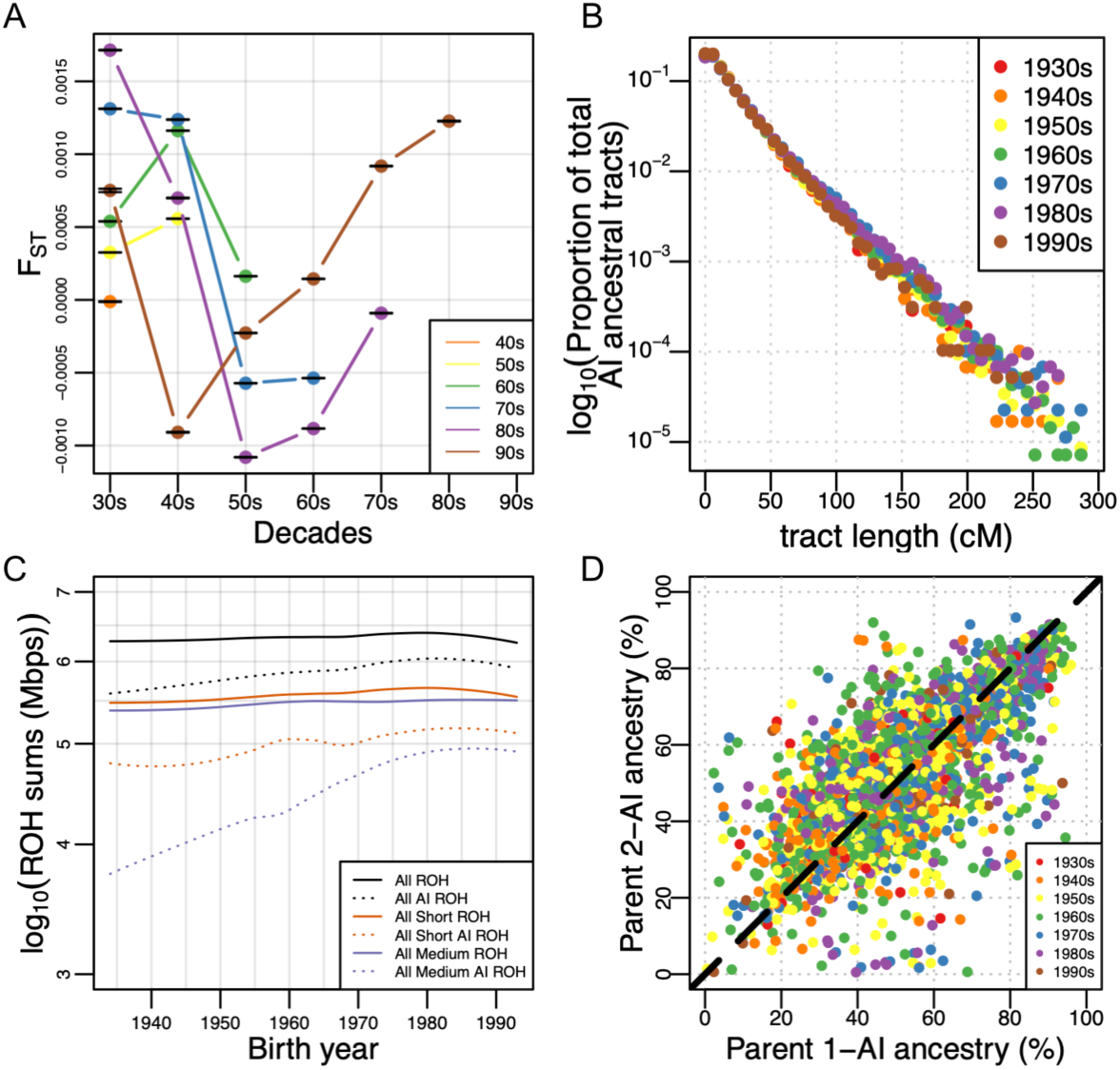
Diversity of and within Amerindigenous ancestral tracts. A) F_ST_ estimates calculated between each decade group. Bars represent the 95% CI. B) Proportion of total Amerindigenous (AI) ancestral tracts in the HCHS/SOL Mexican American population by decade. C) Loess regression of the log of the sum of total ROH and ROH overlapping Amerindigenous (AI) ancestral tracts separated by ROH class. Total long ROH is not represented as an individual group due to the high number of individuals missing long ROH (1694 for long ROH across ancestries and 1987 for long AI ROH) but was included in the sum of “All ROH” and “All AI ROH”. D) Correlation of parent’s inferred global Amerindigenous (AI) ancestries using ANCESTOR.

### Little evidence that Amerindigenous ancestry tract lengths have changed

We next sought to test whether differences at the local ancestry level could explain the shift in global Amerindigenous ancestry over time in the Mexican Americans. We calculated the length of each RFMix inferred local ancestry tract in each Mexican American individual, and tested for differences in the distribution of tract lengths across birth-decades using a multiple linear regression model (see methods). We found no significant associations between the decade bin and the proportion of Amerindigenous ancestral tracts at various lengths (Figure 2B), even when testing for violations of model assumptions (e.g. normalizing the tracts per bin by the number of individuals, or excluding the 1930s and/or 1990s individuals due to the small sample size in each bin).

### Increased runs of homozygosity over time

Since genetic diversity has decreased over time in the Amerindigenous ancestry tracts of Mexican Americans (despite rapid growth of the census population size), it is possible that this population has also experienced increased haplotype homozygosity over time. We investigated this possibility by exploring runs of homozygosity (ROH) in Amerindigenous ancestry tracts in each of the 3622 Mexican Americans. We classified ROH into three categories: short, medium, and long, based on the length distribution in the population. Generally, short ROH are tens of kilobases in length and likely reflect the homozygosity of old haplotypes; medium ROH are hundreds of kilobases in length and likely reflect background relatedness in the population; and long ROH are hundreds of kilobases to several megabases in length and are likely the result of recent parental relatedness. Figure 2C shows a fitted loess curve to the log of the total length of ROH summed across each Mexican American’s genome as a function of their birth year, broken down by ROH size class (as well as the total of each size class that overlaps all ancestry tracts (Supplementary Figure 8A) and Amerindigenous ancestry tracts (Supplementary Figure 8B)). Overall, we find a significant positive correlation between birth year and the total summed ROH across size classes (*τ*=0.0449, P=6.12e-5, Kendall’s rank correlation), but this becomes more significant when we restrict our analysis to ROH calls that overlap Amerindigenous ancestry tracts (*τ*=0.0873, P=9.46e-15). When stratified by size class, the associations (all Kendall’s rank correlation) in ROH were primarily driven by the short (*τ*=0.0833, P=9.45e-14), and medium (*τ*=0.0718, P=1.46e-10) size classes, and are again strongest when ROH overlap Amerindigenous ancestry tracts (short *τ*=0.107, P<2.2e-16, medium *τ*=0.1003, P<2.2e-16). The long ROH had a negative correlation with birth year, but was insignificant after multiple testing (*τ*=−0.0291, P=0.01499; note that 1694 individuals did not have any long ROH calls in their genome).

### Strong ancestry-related assortative mating in HCHS/SOL Mexicans

Given that short and medium length ROH have increased over time, it appears that background relatedness within Amerindigenous ancestry in Mexican Americans has increased over time (but not an increase in recent parental relatedness). One way for this to occur is if individuals with similar ancestry patterns tend to mate with one another more often than expected under a model of random mating (i.e. assortative mating). To measure assortative mating, we estimated the ancestral proportions of the biological parents of each HCHS/SOL Mexican American (see methods). With individuals from all decades pooled together, we found the inferred biological parental Amerindigenous ancestries to be significantly correlated (Figure 2D, r=0.708, 95% CI:0.69-0.72, P<2.2e-16, Pearson correlation). When stratified by decade, the correlation in inferred parental Amerindigenous global ancestry ranged from 0.65 to 0.74 (Supplementary Figure 9), but were not statistically different from each other. This shows that there was a strong parental ancestry correlation among Mexican Americans over different generations. This signature of assortative mating is not due to recent parental relatedness, because there is no trend in long ROH with birth year (and an overall low rate of long ROH among Mexican Americans).

### Genetic correlation of global Amerindigenous ancestry with biomedical traits

We have shown that genetic variation in the Mexican American population is dynamic, with Amerindigenous ancestry increasing over a short period of time (combined with decreased genetic diversity and increased short and medium length ROH within Amerindigenous ancestry tracts). These features may have implications for the genetic architecture of complex traits within Mexican Americans, a topic that is understudied and poorly understood. To further our understanding of the genetic architecture of complex traits in Mexican Americans, we investigated the relationship between Amerindigenous ancestry and various complex traits that may be relevant to biomedical phenotypes. Specifically, we tested for a correlation between 66 complex traits from the HCHS/SOL phenotypic dataset and global Amerindigenous ancestry (Kendall’s *τ*). As illustrated in Figure 3, 22 of these traits (33%) are significantly correlated after Bonferroni correction (P<0.00076). We found that the effect of global Amerindigenous ancestry on many of these phenotypes persisted when using multiple regression to account for age, sex, center, and the sampling weight (Supplementary Table 2), highlighting the need for increased investigation into the role of Amerindigenous genetic ancestry in admixed populations such as Mexican Americans.

**Figure 3.**
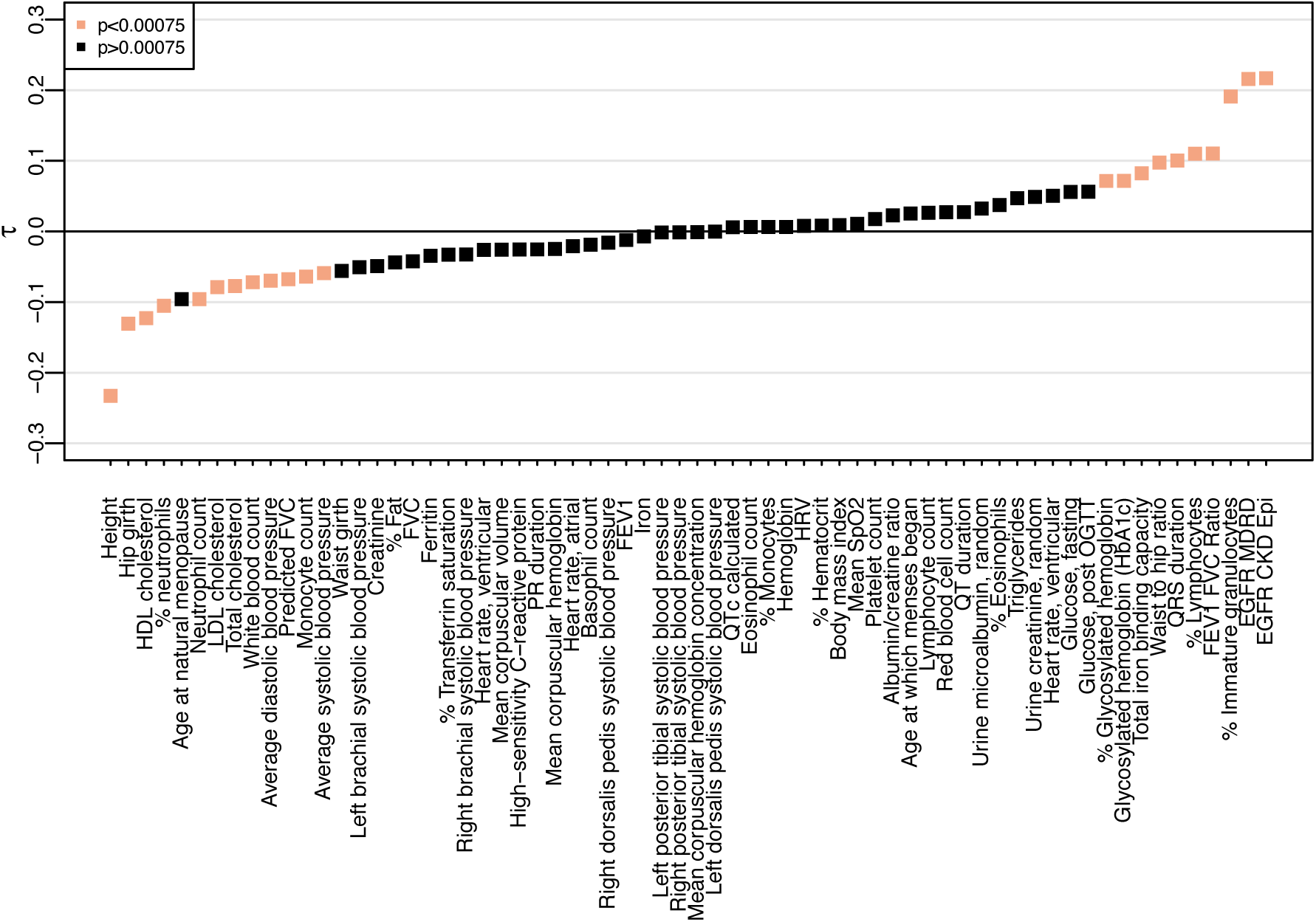
Correlation of 66 quantitative traits with global Amerindigenous ancestry. Significance level was determined using Bonferroni correction adjusting by the number of quantitative traits tested (0.05/66=0.00075).

### Assessing the genetic contribution of Amerindigenous ancestry to height

Among the traits we tested for a correlation with global Amerindigenous ancestry, height had the strongest negative correlation, and our regression model indicated that height also had a strong positive relationship with birth year (Figure 4A and Supplementary Table 3). Globally, populations have grown taller over time due to a variety of non-genetic, environmental factors (26). We find a similar trend in the HCHS/SOL Mexican Americans (Figure 4A). Indeed, when we stratified individuals by quartiles of global Amerindigenous ancestry, we see that all quartiles have increased in height by a similar amount over the period investigated (though individuals with lower Amerindigenous ancestry were taller on average).

**Figure 4:**
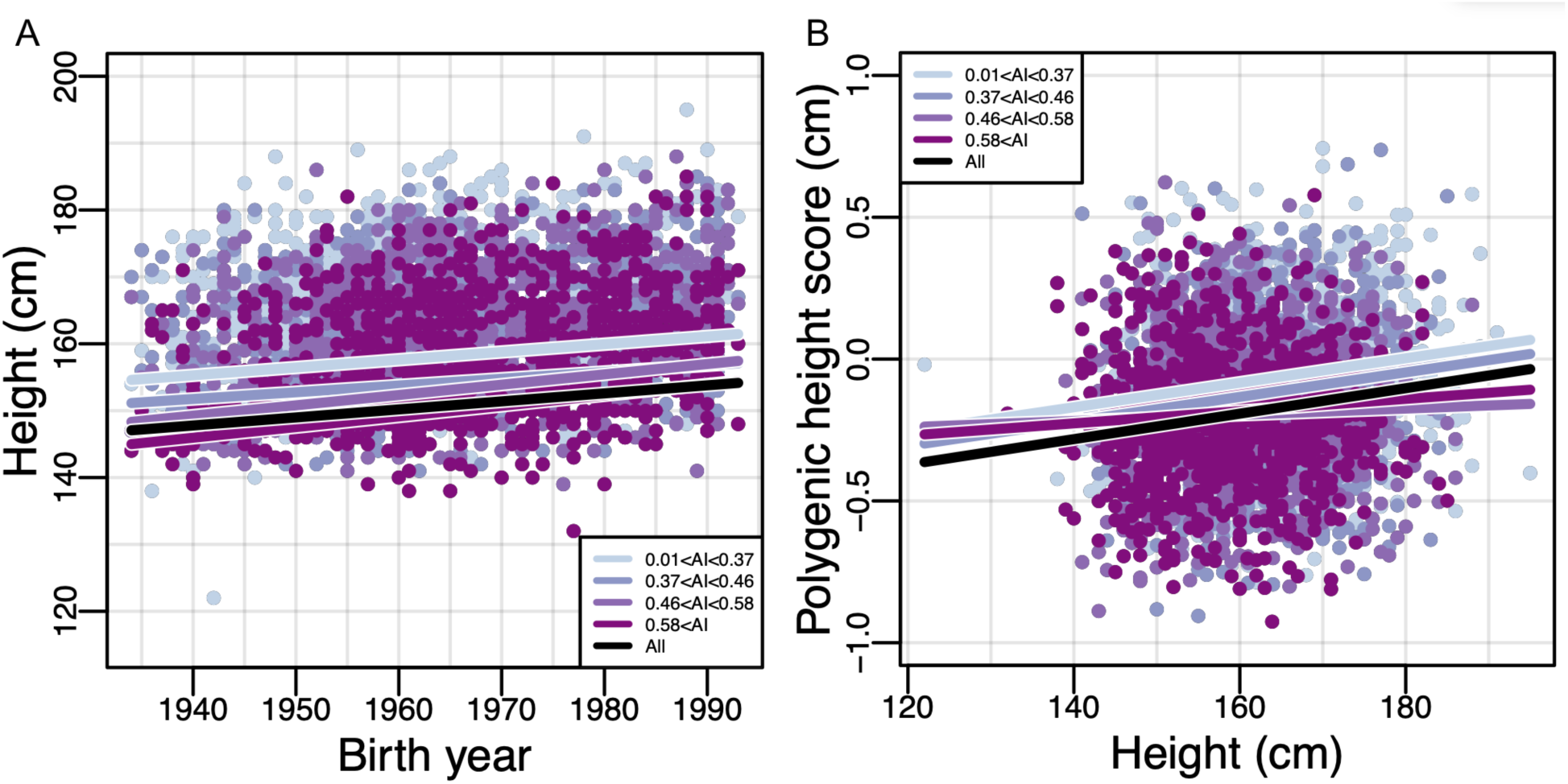
Height and global Amerindigenous ancestry in HCHS/SOL Mexican Americans. Each plot illustrates the relationship between A) Birth year and height B) Height and polygenic height score (PHS). The black line indicates the fitted linear model for all individuals. Each color represents a different quartile of Amerindigenous global ancestry. Polygenic height scores were assessed utilizing UKBB summary statistics for 1,128 SNPs.

Height is one of the most highly studied complex traits, with GWAS sample sizes numbering in the hundreds of thousands (27). Results for many of these studies have been made readily available on public databases as summary association statistics that can be leveraged to build genetic predictions through polygenic risk scores (PRS) (28). In Europeans, PRS have been shown to have great predictive power for several traits, including breast cancer, prostate cancer, and type 1 diabetes (22, 29–31). PRS are most effective in populations of European descent as GWAS studies have been primarily performed in these populations (21–23) and are expected to be biased when applied to other populations due to differences in the genetic architecture of traits across diverse populations (32). Since Mexican Americans have some fraction of European ancestry, we sought to determine whether PRS calculated utilizing GWAS summary statistics from European populations could still provide useful insight.

To evaluate the effectiveness of PRS for height (i.e. the polygenic height score, or PHS, see methods), we first tested whether there was an association between the observed height and the predicted height estimates while controlling for sampling weight, sex, and recruitment center (see methods). We identified a significant association between observed height and predicted height for the population as a whole (*β*=0.0044881, P=2.19e-12; Figure 4B, Supplementary Table 4). However, when we stratified by quartiles of Amerindigenous global ancestry, the association only remained for the individuals in the lower two quartiles of global Amerindigenous ancestry proportions (AIA<0.37: *β*=0.004, P=0.0008 and 0.36<AIA<0.46: *β*=0.004, P=0.003, Supplementary Table 4). The association between predicted height and observed height was no longer significant for individuals in the upper two quartiles of global Amerindigenous ancestry proportions (0.46<AIA<0.58: *β*=0.0011, P=0.39 and 0.58<AIA: *β*=0.0022, P=0.08, Supplementary Table 4).

As we had found global Amerindigenous ancestry to be increasing over time, we hypothesized that there would be a change in PHS over time as well. However, we find little evidence supporting this hypothesis. While individuals born earlier than 1950 or in the 1950s have a stronger correlation between their PHS and observed height (*β*=0.034 and 0.039; p=5.6e-4 and 2.7e-7 respectively) than individuals born in the 1960s, 1970s, or 1980s (*β*=0.016, 0.029, and 0.029; p=0.044, 0.0066, and 7.8e-5 respectively), there is no clear trend and we did not find a significant effect of birth year on PHS (P=0.09) even when we stratified by the quartiles of global Amerindigenous ancestry.

## Discussion

The United States is a dynamic, rapidly changing population, and this will continue to occur as the population size grows (20). Hispanics/Latinos are the largest and fastest growing minority group, and are projected to comprise over 25% of the US population by 2060. They are a genetically and phenotypically diverse population as a result of extensive admixture between Amerindigenous populations and immigrants from multiple geographic locations around the world. In this study, we identified additional population substructure complexities that may contribute to phenotypic variation within Hispanics/Latinos.

Specifically, we demonstrated how the admixture dynamics of Mexican Americans have changed over time, resulting in an increase of ∼20% Amerindigenous ancestry on average over the 50-year period studied. This change in ancestry is equivalent to a mean increase in Amerindigenous ancestry of ∼0.4% per year. While the effect sizes vary to some extent, we replicate the underlying pattern across multiple data stratifications (two metropolitan cities, US born and non-US born) and also replicate this feature in an independent cohort of Mexican Americans. Further, we find that a similar trend holds across multiple self-identified Hispanic/Latino populations in the US (and is statistically significant in Central Americans). This effect does not appear to have a simple explanation: we do not see any statistically significant increases at individual loci, we do not see more than a negligible degree of population differentiation over time, and this increase cannot be entirely explained by very recent migration. We do, however, find that as Amerindigenous ancestry has increased, genetic diversity within Amerindigenous ancestry tracts across Mexican Americans has decreased over time, and is associated with increased short and medium length ROH over time. This suggested that there could be increased relatedness within Amerindigenous ancestries within Mexican Americans, and we confirmed that there is a very high degree of ancestry-based assortative mating within the Mexican American population.

What could be driving the increased Amerindigenous ancestry in Mexican Americans? Population genetic theory suggests that while assortative mating could result in increased ROH and decreased genetic diversity, ancestry-based assortative mating alone should not result in mean changes in global ancestry proportions. Regardless of the underlying mechanisms driving increased Amerindigenous ancestry in Mexican Americans, this additional source of temporal substructure within this population has substantial consequences for phenotypic variation in biomedical traits. We identify several biomedical traits that are correlated with Amerindigenous ancestry, and show that in the case of height, there are both ancestry and temporal effects. Further study is necessary to understand whether other biomedical traits are also changing over time as the genomic ancestry proportions change in this population.

Interestingly, we identified another source of structure within HCHS/SOL, particularly in the African ancestral component of Hispanics/Latinos. In our UMAP analysis, the YRI sample form their own cluster as a reference population as compared to the Amerindigenous and European reference populations which border the admixed samples with the highest proportion of each ancestry, respectively (Supplementary Figure 1C). While most Latin Americans can trace their African ancestry to Sub-Saharan Africa, previous studies have also identified hidden Northern African ancestry in individuals from Southern Europe (33–35), who primarily colonised the Americas. This may explain why the YRI sample is not at the boundary of the individuals with the highest proportion of African ancestry in the HCHS/SOL sample as the African ancestral component may be more complex. Our results suggest how careful consideration must be taken into account when selecting reference populations to study the African ancestral component of admixed individuals from the Americas.

In our study, we bring specific attention to the biases that continue to exist with using European GWAS summary statistics to calculate polygenic risk scores in admixed populations such as Mexican Americans that are comprised of European, Amerindigenous, and African genetic ancestries. In particular, in the case of height, we found that the polygenic height score (PHS) correlated with observed height only in the subset of individuals with the lowest levels of Amerindigenous ancestry (i.e. the subset of individuals with highest European ancestry). As the population dynamics of the US continue to change, it is imperative that we study diverse populations, or we risk exacerbating the health disparities that currently exist. To date, population-based medical genomics research (and its subsequent benefits) have been disproportionately focused on populations of European ancestry. In order to improve the design and implementation of medical genetics studies for the ethnically diverse U.S. population, we need detailed insights into the population history of diverse U.S. populations. This includes characterizing the admixture dynamics of Hispanic/Latino populations, as well as the evolutionary forces that shaped patterns of genetic variation of the ancestral populations that contributed to modern day Hispanic/Latino populations.

The genetic variation of the Hispanic community in the United States belies categorization under a single label (10). The events that have shaped and continue to shape this genetic diversity are complex, numerous, and nuanced, and the social history of such a diverse population is intrinsic to any genetic study. Mexico’s society was largely defined by an established social caste system based on ancestry, which disappeared after Mexico’s independence in 1821 (36). Even so, social inequalities persist today with skin colour having a significant effect on wealth and education (37). A multitude of factors within and outside Mexico — whether related to trade, immigration policies, or armed conflicts — acted to influence who immigrated to the United States, and the impact of each of these fluctuates over time (38–40). These changes shift the demographics of immigration, which is inherently related to the genetic ancestry of the population.

Consequently, this shapes the genetic architecture of complex traits. Diverse populations are at risk not only from underrepresentation in research, but because of poor understanding of the temporal and spatial dynamics at play in genetic variation. The promise of equitable precision medicine — one of the ultimate goals of medical genomics — cannot be kept without understanding this interplay. Health disparities in the United States are fed by structural inequalities. For example, studies that use modern Artificial Intelligence techniques have already been shown to inflate existing disparities between Black Americans and White Americans (41). Such biases, whether from algorithms, study designs, or misunderstandings of subtleties in data, feed into the larger systemic pressures faced by minority populations in the United States.

While we have shown a dramatic shift in ancestry proportions in US Hispanic/Latinos, one of the caveats of this study is that the HCHS/SOL cohort is not representative of all US Hispanics/Latinos. HCHS/SOL participants were recruited at four primary centers: Bronx, Chicago, Miami, and San Diego. There may be additional genetic diversity that has not been captured by this dataset and trends exhibited in this dataset may not translate to Hispanic/Latino populations living in other regions of the US (though the temporal increase in Amerindigenous ancestry was replicated in an independent sample of Mexican Americans). Further, we have only assembled a reference panel with limited numbers of individuals with various Amerindigenous, European, and African ancestry. With better population genetic modeling and a deeper understanding of the social and historical aspects of Hispanic/Latino populations, we will be able to improve our understanding of the genetic and phenotypic diversity across these populations, and subsequently improve our ability to understand genetic contributions to complex traits and disease. These insights will lead to optimization of population sampling for the design of future medical genetic studies, the identification of disease risk variants, and ultimately, precision medicine for all.

## Methods

### Study dataset and initial quality control

The HCHS/SOL study is a community-based cohort study of self-identified Hispanic/Latino individuals from four US metropolitan areas with the general goal of identifying risk and protective factors for various medical conditions including cardiovascular disease, diabetes, pulmonary disease, and sleep disorders (25). 12,434 participants with birth year estimates between 1934-1993 who self-identified as being of Cuban, Dominican, Puerto Rican, Mexican, Central American, or South American background consented to genetics studies and posting of their genetic and phenotype data on the publicly available Database of Genotypes and Phenotypes (dbGaP) through Study Accession phs000810.v1.p1. Samples were genotyped on an Illumina custom array, SoL HCHS Custom 15041502 array (annotation B3, genome build 37), consisting of the Illumina Omni 2.5M array and 148,353 custom single nucleotide polymorphisms (SNPs) (10). Data posted to dbGaP had passed initial sample quality control filters, including removing samples with differences in reported vs. genetic sex, call rates > 95%, and evidence for sample contamination (e.g. heterozygosity and sample call rates). For initial SNP quality control, we filtered out SNPs that were monomorphic, positional duplicates, or Illumina technical failures, as well as SNPs that had cluster separation <= 0.3, call rate <=2%, >2 disconcordant calls in 291 duplicate samples, >3 Mendelian errors in parent-offspring pairs/trios, Hardy-Weinberg Equilibrium combined P-value <10^−5^, and sex differences in allele frequency ≥0.2. Our filtering resulted in 1,763,935 genotyped SNPs with minor allele frequency (MAF) >0.01.

Additional sample quality control performed in the HCHS/SOL dataset included filtering out samples with 1) large chromosomal anomalies, 2) substantial Asian ancestry as previously identified in HCHS/SOL (*12*) and 3) individuals with up to third degree genetic relatedness in the dataset as inferred by REAP (42). For genetic relatedness filtering, individuals from pairs were kept to maximize representation of the birth year distribution, which resulted in 10,268 unrelated remaining individuals.

From the original HCHS/SOL analysis, individuals were classified into genetic-analysis groups, similar to self-identified background groups in that they share cultural and environmental characteristics, but are also more genetically homogenous (10).

Birth year for all individuals was estimated by subtracting the difference between date of first clinic visit for the baseline examination (25) and age. Year of arrival was estimated by subtracting the difference between date of first clinic visit for the baseline examination and years in the US.

### Global, local, and parental ancestry inference

All ancestry analyses were restricted to the 211,152 autosomal SNP markers that overlapped between the study and reference panel genotyping array. For the HCHS/SOL dataset, global African, European, and Amerindigenous ancestries were inferred with ADMIXTURE, in an unsupervised manner, with K=3. Amerindigenous ancestry refers to estimates of Indigenous genetic ancestry from the Americas. For some analyses, HCHS/SOL individuals with greater than 95% of a single ancestry (e.g African, European, or Amerindigenous) were filtered out resulting in 9,913 individuals: 1,099 Central American, 1,536 Cuban, 954 Dominican, 3,622 Mexican, 1,783 Puerto Rican, 652 South American and 267 “Other” individuals.

Ancestral tracts, known as ‘local’ ancestry, along the genome for all HCHS/SOL individuals were inferred using RFMix (43) and a three population reference panel, comprised of 315 individuals: 104 HapMap phase 3 CEU (European) and 107 YRI (African) individuals (44) and 112 Amerindigenous individuals from throughout Latin America (8). The reference panel was limited to individuals with 99% continental ancestry as inferred by unsupervised ADMIXTURE (45). Prior to local ancestry inference, HCHS/SOL individuals were merged with the reference panels and then phased using SHAPEIT2 (46). For all HCHS/SOL Mexican American individuals, parental genomic ancestry was inferred with ANCESTOR (47) using the local ancestry estimates generated by RFMix.

### Uniform Manifold Approximation and Projection (UMAP)

Principal components for HCHS/SOL and the reference panel were computed using smartPCA (48). UMAP (version 0.3.8) was run using the Python script freely available at https://github.com/diazale/gt-dimred with parameter specification set at 15 nearest neighbours and a minimum distance between points of 0.5.

### Admixture mapping

Local ancestry estimates for 211,151 SNPs across the genome were used to perform admixture mapping in HCHS/SOL Mexican Americans to determine if younger individuals harbored excess Amerindigenous ancestry in certain regions of the genome. Admixture mapping was performed applying two different models: 1) a linear regression model with age as the dependent variable adjusting for global Amerindigenous ancestry, sampling weight and center and 2) a logistic regression model dividing the HCHS/SOL Mexican cohort in to an older vs younger generation with 1965 set as the dividing point while also adjusting for global Amerindigenous ancestry, sampling weight, and center. The threshold for genome-wide significance, 1.38×10^−4^ was calculated using the empirical autoregression framework with the package *coda* in R to estimate the total number of ancestral blocks (49, 50).

### Tract Lengths

The multiple regression model: log(*f*) =*β*_0_ +*β*_1_ *T* +*β*_2_ *A* +*β*_3_ *TA* +*ε*, where *f* is a matrix containing the proportion of lengths of all ancestral tracts across the genome for all 3622 Mexican American individuals, *T* the tract length bin and *A* decade of birth year bin, was used to test for an effect of birth decade on the proportion of Amerindigenous ancestral tract lengths. For assessment between the fraction of ancestry tracts in an individual’s genome and birth year, long tract cutoffs were chosen based on tract separation between the birth year decades in Figure 2B.

### Diversity Calculations

Subcontinental ancestry was assessed using the diversity measurements π and F_ST_. π was calculated as the average number of pairwise genetic differences among all pairs of overlapping Amerindigenous ancestry tracts across individuals. F_ST_ was calculated as:

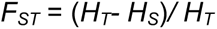

where *H_T_* is the average heterozygosity when all individuals are pooled across decades and *H_S_* is the average heterozygosity within each decade of individuals.

### Inference of Runs of Homozygosity

Runs of homozygosity (ROH) were called using the program GARLIC v1.1.4 (51) on 211,152 sites for the Mexican American individuals. An analysis window size of 50 SNPs and an overlap fraction of 0.25 were both chosen using GARLIC’s rule of thumb parameter estimation. GARLIC chose a LOD score cutoff of 0. Using a three-component Gaussian mixture, GARLIC determined class A/B (short/medium) and class B/C (medium/long) size boundaries as 845,097 bp and 2,501,750 bp, respectively.

### Imputation

Imputation for HCHS/SOL was performed locally using IMPUTE2 with the 1000 Genomes Project Phase 3 haplotypes used as a reference panel. After filtering on an info score cutoff of 0.3, this resulted in 33,041,084 SNPs.

### Polygenic Risk Score Calculations

Polygenic risk scores for height were calculated using the publicly available UK Biobank (UKBB) GWAS Round 2 Summary Statistics retrieved from http://www.nealelab.is/uk-biobank. Briefly, for sample quality control, sample inclusion was limited to unrelated samples who passed the sex chromosome aneuploidy filter. British ancestry was determined using the 1^st^ 6 PCs; individuals more than 7 standard deviations away from the 1^st^ 6 PCs were excluded. Further filtering included limiting to self-reported ‘white-British’ / ‘Irish’ / ‘White’ resulting in a QCed sample count of 361,194 individuals https://github.com/Nealelab/UK_Biobank_GWAS#imputed-v3-sample-qc. An imputation panel of ∼90 million SNPs from HRC, UK10K and 1KG were used to impute genotypes. 13.7 million autosomal and X-chromosome SNPs passed quality control thresholds including Info score>0.8, MAF>0.0001, and HWE p-value>1e-10. For the phenotype, a linear regression model in Hail was run for all individuals (both sexes) adjusting by the first 20 PCs + sex + age + age^2^ + (sex*age) + (sex*age)^2^. For height, there was complete phenotype information for 360,388 individuals.

Risk scores were calculated by extracting the overlapping genome-wide significant hits initially discovered in the UKBB GWASs of height and selecting SNPs with the lowest p-value in each 1Mb window across the genome. For height this resulted in a dataset of 1,103 overlapping SNPs that were present in our dataset of genotyped and imputed SNPs.

### Health and Retirement Study (HRS)

For replication, we used genotype data from 705 self-identified Mexican-Americans from the Health and Retirement Study (HRS) (52), genotyped on the Illumina Human Omni 2.5M platform. HRS data was made available under IRB Study No. A11-E91-13B - The apportionment of genetic diversity within the United States. Estimated global ancestry proportions for the Mexican American population in the HRS were calculated as in Baharian et al. (12), which used an alternative reference panel and alternative ancestry inference approach. Briefly, RFMix was used to infer local ancestry estimates across the genome utilizing CHS, YRI, and CEU individuals from the 1000 Genomes Project as reference populations for Amerindigenous/Asian, African, and European ancestries, respectively. Global ancestry estimates were calculated using the summed RFMix calls.

### Statistical Analyses and Plots

Statistical analyses and plot generation were performed within Rstudio using Version 1.1.463 and R version 3.5.3. ternary and ggridges/ggplot2 packages were used to create the simplex and ridgeline plots.

## Acknowledgements

We thank many colleagues who commented on our preprint prior to submission, particularly Reed Cartwright for suggestions on terminology. MLS was supported through the National Human Genome Research Institute (NHGRI) of the National Institutes of Health (NIH) under Award Number F31HG010104. ADP and SG were supported, in part, thanks to funding from the Canada Research Chairs program and CIHR grant MOP-136855. RDH was supported, in part, by NHGRI grant R01HG007644 and the Canadian Research Chairs program.

## Supplementary Material

**Supplementary Figure 1.**
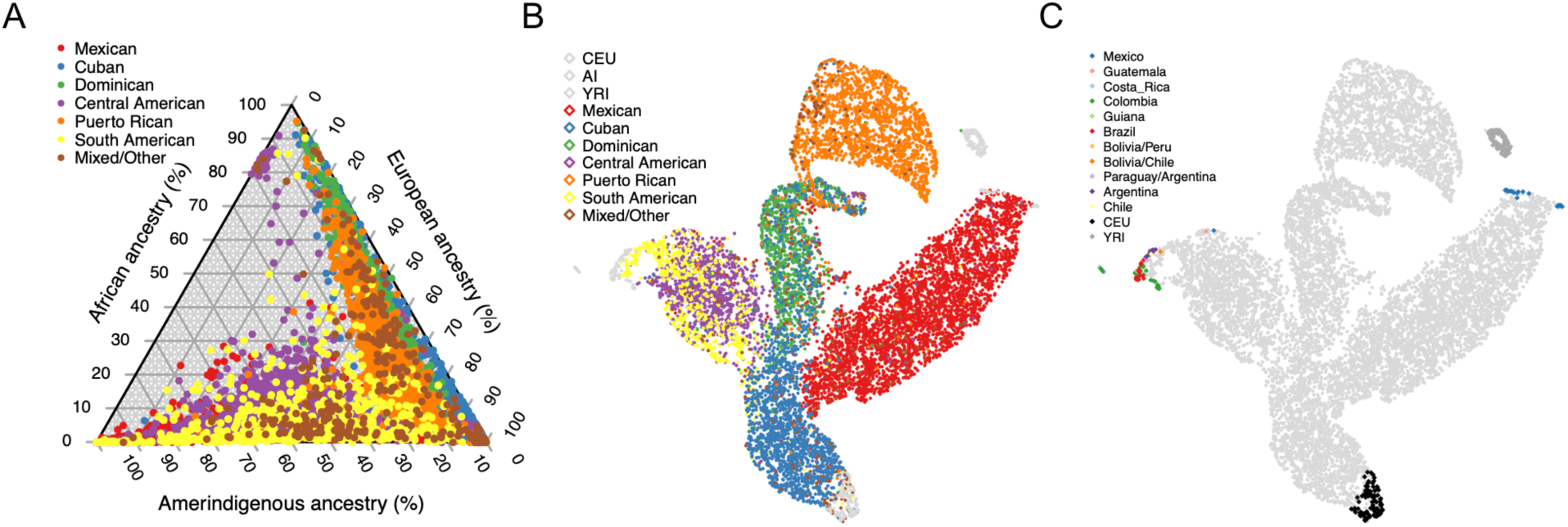
Continental ancestral diversity of HCHS/SOL. A) Ternary plot of global ancestry proportions colored by population for 10,268 HCHS/SOL individuals B) Uniform Manifold Approximation and Projection (UMAP) plot of HCHS/SOL and the reference panel (n=10,591) using 3 principal components, colored by HCHS/SOL population. C) UMAP plot of HCHS/SOL and the reference panel (n=10,591) using 3 principal components, colored by reference population.

**Supplementary Figure 2.**
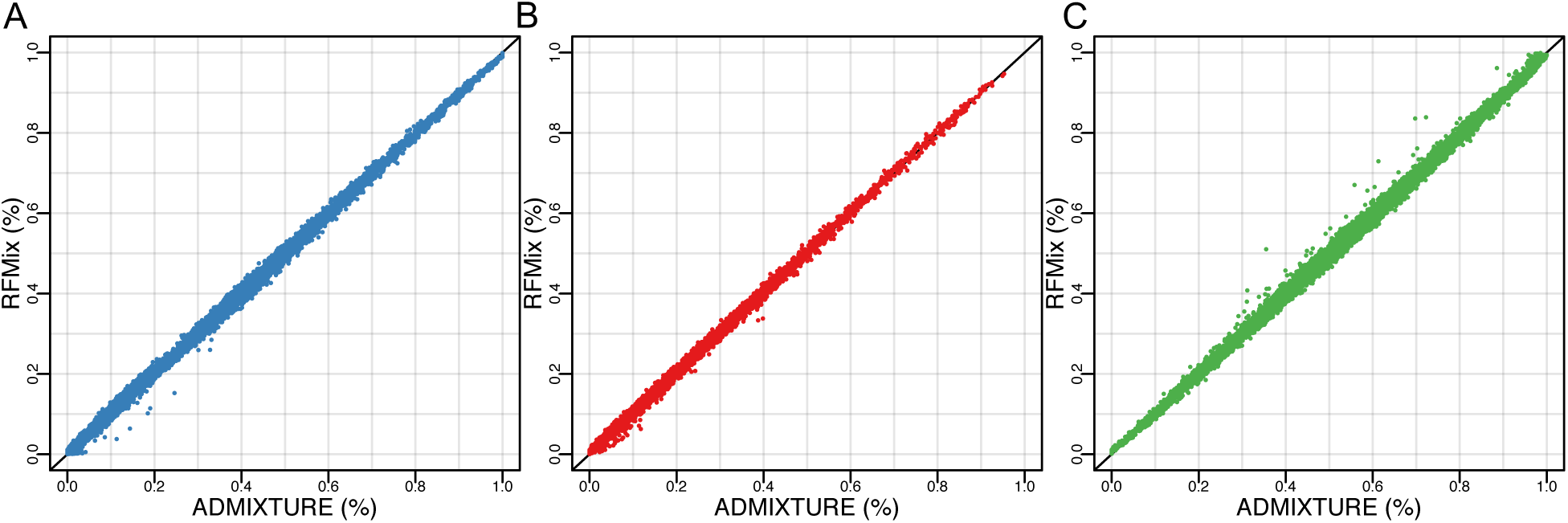
Concordance of ADMIXTURE and RFMix global ancestry estimates. A) Amerindigenous ancestry B) African ancestry and C) European ancestry.

**Supplementary Figure 3.**
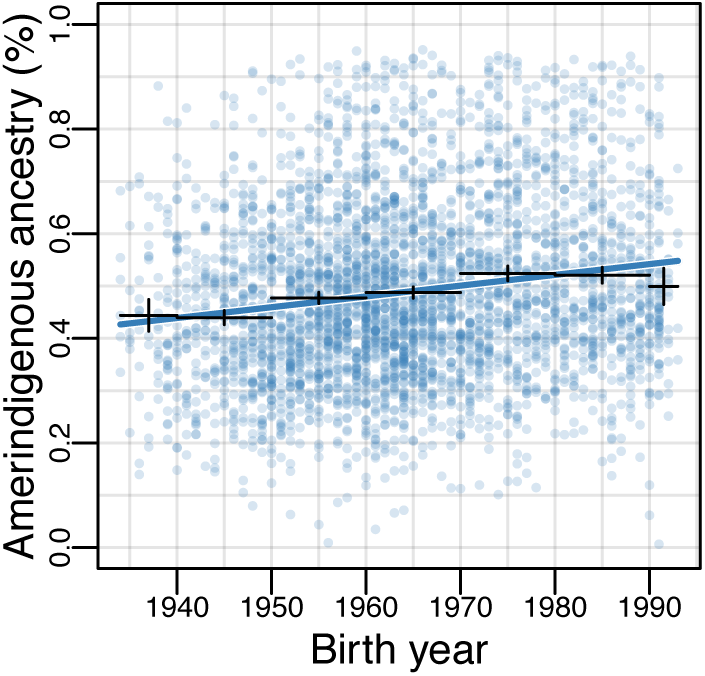
RFMix inferred Amerindigenous global ancestry proportions plotted over time for HCHS/SOL Mexican Americans (n=3622).

**Supplementary Figure 4:**
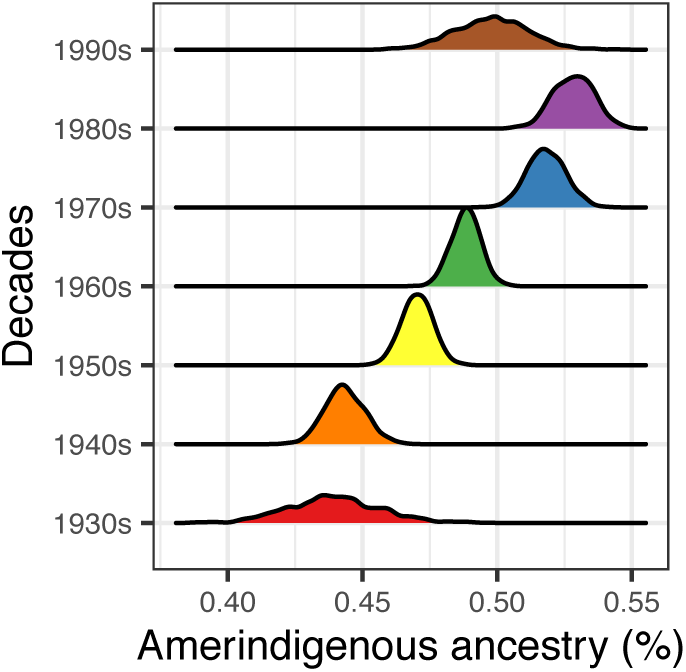
Distributions of Amerindigenous global ancestry means for HCHS/SOL Mexican Americans (n=3622) generated by 1000 bootstrap resampling iterations within each decade of binned birth years.

**Supplementary Figure 5.**
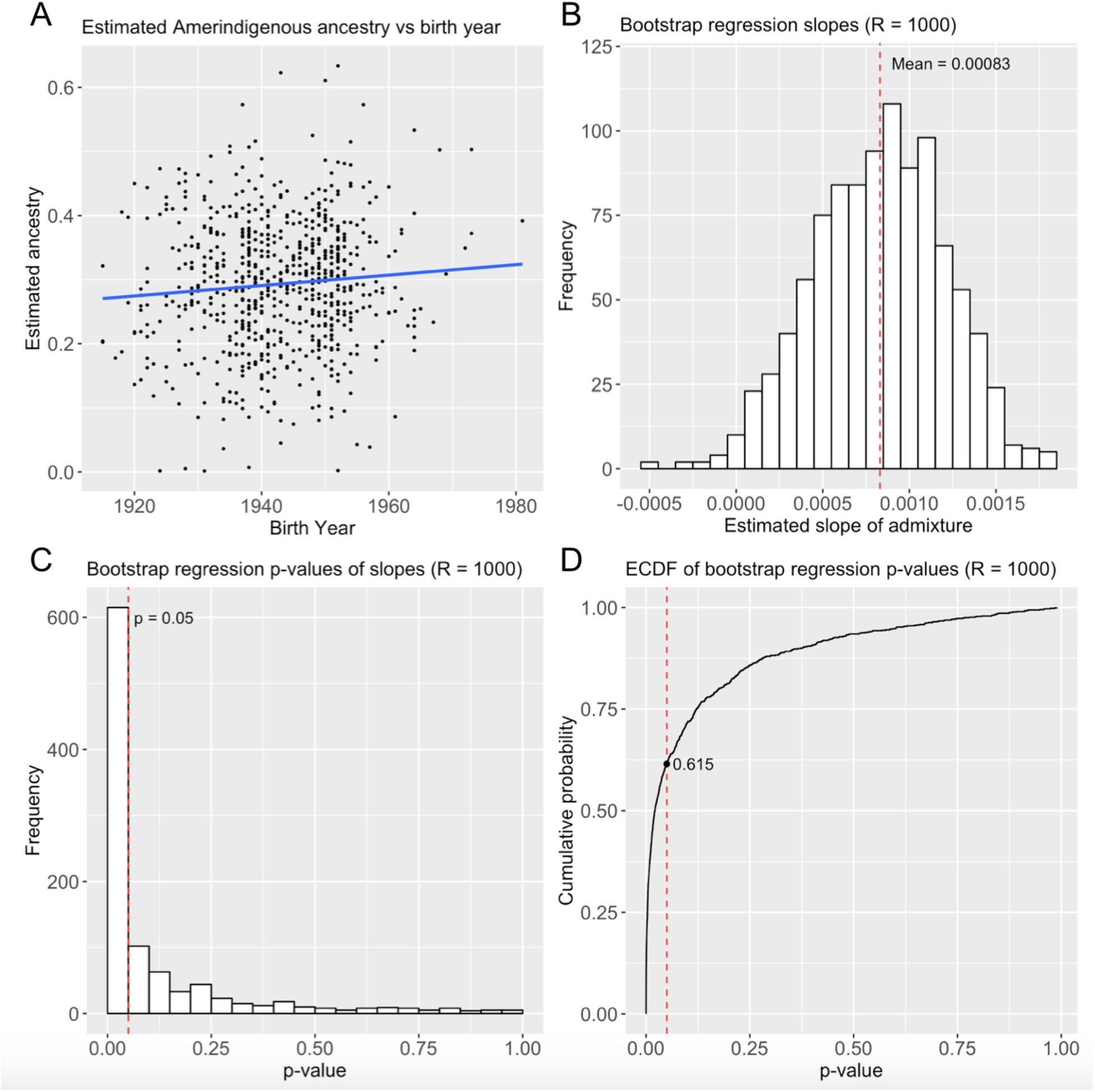
Replication in the Health and Retirement Study for 705 self-identified Mexican Americans. A) Ancestry over time B) Distribution of regression slopes after 1000 bootstrap resampling iterations C) Distribution of bootstrap regression p-values D) ECDF of bootstrap regression p-values.

**Supplementary Figure 6.**
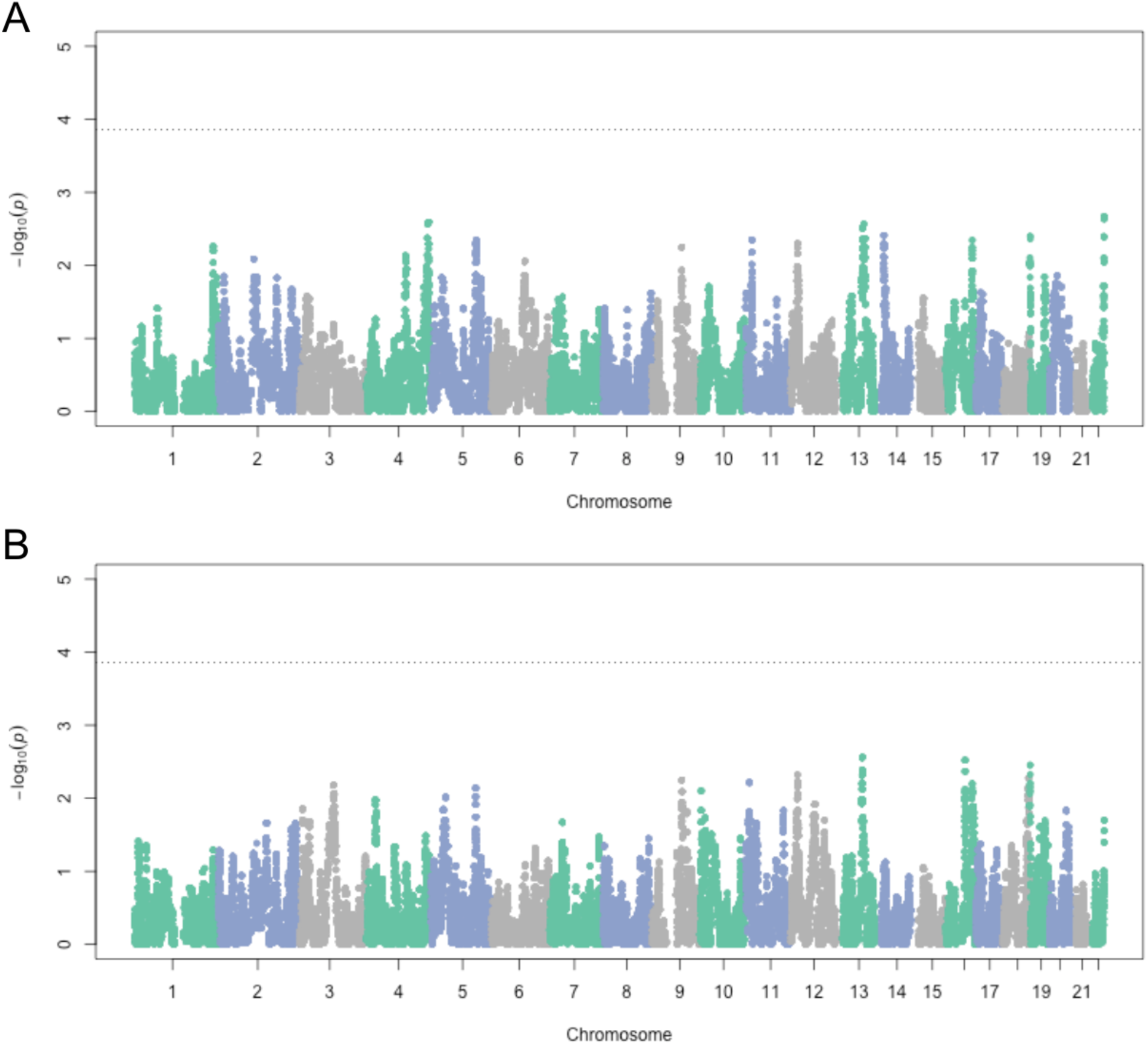
Admixture mapping in HCHS/SOL Mexicans (n=3622) for Amerindigenous ancestry and A) birth year and B) generation. Ancestry association testing was performed at 211,151 markers using A) linear regression and B) logistic regression, both including global Amerindigenous ancestry, sampling weight and center as covariates.

**Supplementary Figure 7.**
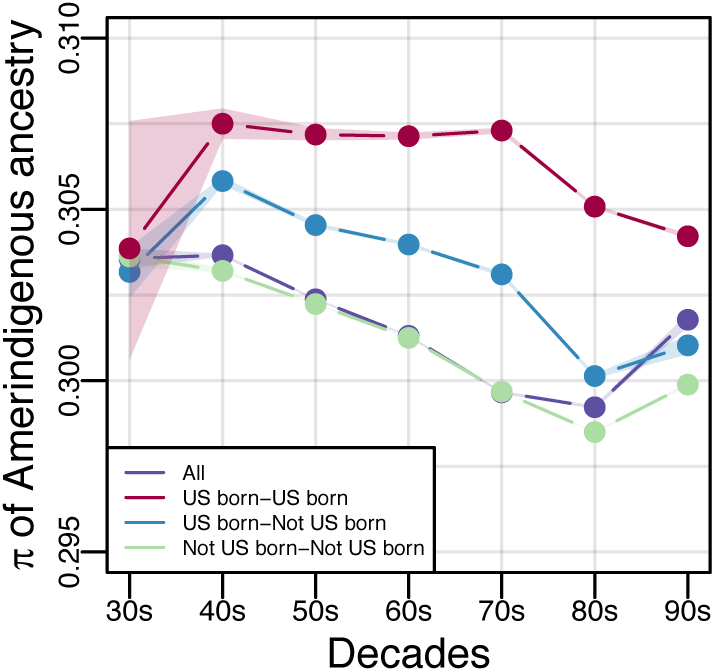
Diversity of and within Amerindigenous ancestral tracts. Diversity (π) of subcontinental Amerindigenous ancestry stratified by US born/not US born status. π was calculated between pairs within each decade of birth years. 95% confidence intervals are highlighted by the shaded regions for each group.

**Supplementary Figure 8.**
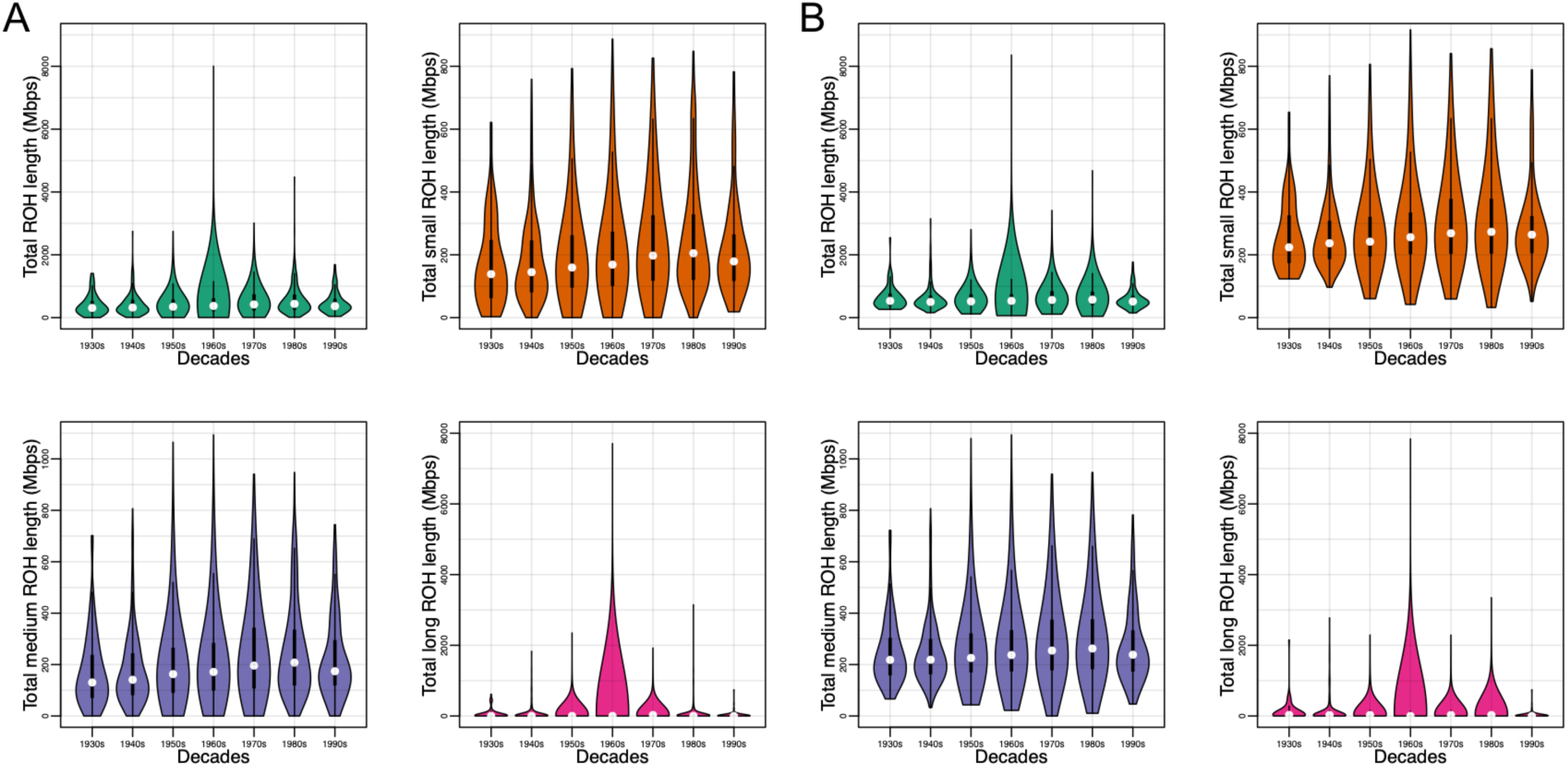
Runs of homozygosity (ROH) in HCHS/SOL Mexican Americans. A) ROH across all ancestries separated by ROH class B) ROH overlapping Amerindigenous haplotypes separated by ROH class.

**Supplementary Figure 9.**
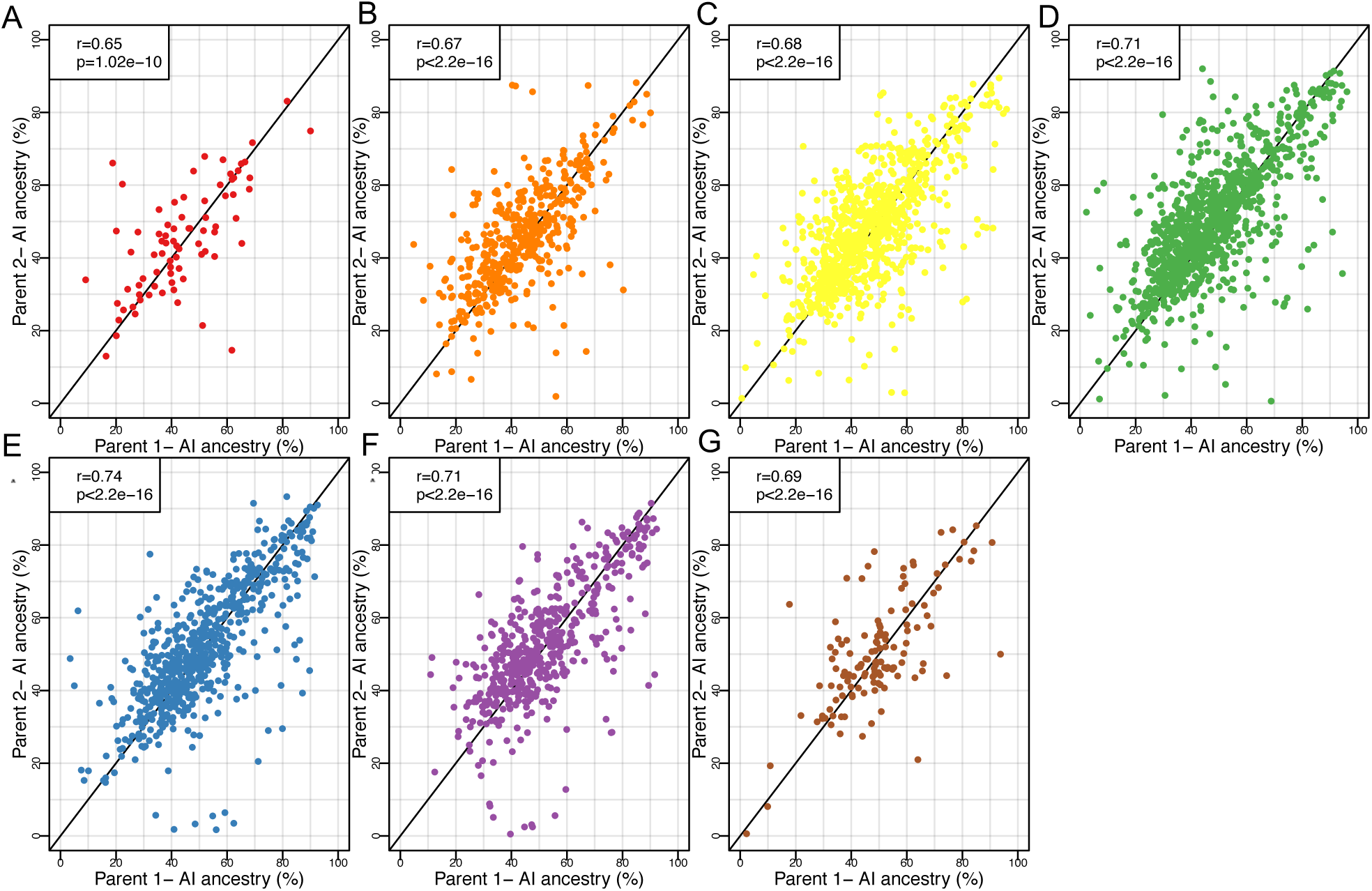
Ancestry-related assortative mating in HCHS/SOL Mexican Americans separated by decade. Each plot represents the correlation of parent’s inferred Amerindigenous (AI) ancestries using ANCESTOR by decade beginning with the 1930s (A) and ending with the 1990s (G). Each point corresponds to one Mexican American couple and the axes correspond to the inferred Amerindigenous (AI) ancestry of each partner.

**Supplementary Table 1:** Association of global ancestries and birth year for all HCHS/SOL individuals. For each population, we tested for an association between global ancestry and birth year while accounting for the sampling design. AI, AFR, and EUR refer to Amerindigenous, African, and European ancestry respectively.

| Population | Ancestry | N | R2 | Effect | Std.Err | P |
| --- | --- | --- | --- | --- | --- | --- |
| Central American | AI | 1099 | 0.0139 | 0.0013 | 0.0004 | 0.0013 |
| Central American | AFR | 1099 | 0.0136 | 0.0002 | 0.0004 | 0.6561 |
| Central American | EUR | 1099 | 0.0138 | -0.0015 | 0.0004 | 0.0002 |
| Cuban | AI | 1536 | 0.0023 | 0.0002 | 0.0001 | 0.0938 |
| Cuban | AFR | 1536 | 0.0014 | -0.0005 | 0.0004 | 0.1490 |
| Cuban | EUR | 1536 | 0.0005 | 0.0003 | 0.0004 | 0.3879 |
| Dominican | AI | 954 | 0.0035 | 0.0002 | 0.0001 | 0.0663 |
| Dominican | AFR | 954 | 0.0030 | -0.0007 | 0.0004 | 0.1287 |
| Dominican | EUR | 954 | 0.0022 | 0.0005 | 0.0004 | 0.2374 |
| Mexican | AI | 3622 | 0.0268 | 0.0023 | 0.0002 | 3.58E-22 |
| Mexican | AFR | 3622 | 0.0008 | 0.0000 | 0.0000 | 0.4189 |
| Mexican | EUR | 3622 | 0.0285 | -0.0023 | 0.0002 | 0.0000 |
| Puerto Rican | AI | 1783 | 0.0014 | 0.0001 | 0.0001 | 0.1533 |
| Puerto Rican | AFR | 1783 | 0.0014 | 0.0003 | 0.0002 | 0.1743 |
| Puerto Rican | EUR | 1783 | 0.0027 | -0.0005 | 0.0002 | 0.0355 |
| South American | AI | 652 | 0.0110 | 0.0016 | 0.0007 | 0.0211 |
| South American | AFR | 652 | 0.0027 | -0.0002 | 0.0004 | 0.5053 |
| South American | EUR | 652 | 0.0080 | -0.0014 | 0.0006 | 0.0335 |

**Supplementary Table 2.** Multiple regression table with traits that were significantly correlated with global Amerindigenous ancestry.

| Trait | N | R2 | Effect | Std.Err | P |
| --- | --- | --- | --- | --- | --- |
| Height | 3615 | 0.588 | -13.637 | 0.676 | 1.01E-85 |
| Predicted FVC | 3522 | 0.851 | -695.376 | 36.867 | 1.13E-75 |
| EGFR MDRD | 3308 | 0.196 | 23.398 | 2.435 | 1.39E-21 |
| EGFR CKD Epi | 3308 | 0.441 | 14.752 | 1.581 | 1.90E-20 |
| Waist to hip ratio | 3617 | 0.195 | 0.043 | 0.007 | 6.96E-10 |
| Glycosylated hemoglobin (HbA1c) | 3609 | 0.081 | 9.184 | 1.590 | 8.30E-09 |
| % Glycosylated hemoglobin | 3609 | 0.080 | 0.837 | 0.146 | 9.88E-09 |
| HDL cholesterol | 3621 | 0.095 | -6.740 | 1.372 | 9.39E-07 |
| Total iron binding capacity | 3620 | 0.066 | 23.540 | 5.467 | 1.71E-05 |
| FEV1 FVC Ratio | 3505 | 0.176 | 2.560 | 0.639 | 6.37E-05 |
| % Lymphocytes | 3442 | 0.021 | 3.768 | 1.023 | 2.35E-04 |
| Hip girth | 3617 | 0.077 | -4.452 | 1.273 | 4.78E-04 |
| % Neutrophils | 3442 | 0.039 | -3.649 | 1.178 | 1.96E-03 |
| Monocyte count | 3443 | 0.024 | -0.048 | 0.019 | 1.27E-02 |
| Neutrophil count | 3442 | 0.043 | -0.399 | 0.165 | 1.59E-02 |
| LDL cholesterol | 3529 | 0.047 | -8.269 | 3.963 | 3.70E-02 |
| Average diastolic blood pressure | 3616 | 0.072 | -2.067 | 1.136 | 6.88E-02 |
| Total cholesterol | 3622 | 0.065 | -6.248 | 4.614 | 1.76E-01 |
| White blood cell count | 3442 | 0.032 | -0.260 | 0.208 | 2.12E-01 |
| Average systolic blood pressure | 3619 | 0.211 | 1.590 | 1.712 | 3.53E-01 |
| % Immature granulocytes | 555 | 0.261 | -0.090 | 0.110 | 4.14E-01 |
| QRS duration | 3596 | 0.168 | 0.062 | 1.268 | 9.61E-01 |

**Supplementary Table 3:** Height over time: Height (cm) as a function of birth year adjusting by sex, center, and sampling weight for 3614 Mexican Americans stratified by the quartiles of global Amerindigenous ancestry (AIA).

| Group | N | R2 | Effect | Std.Err | P |
| --- | --- | --- | --- | --- | --- |
| All | 3614 | 0.54186 | 0.1200379 | 0.00905366 | 3.28E-39 |
| AIA>0.58 | 929 | 0.5745454 | 0.158895 | 0.01783895 | 2.73E-18 |
| 0.46<=AIA<=0.58 | 955 | 0.5784636 | 0.1542762 | 0.01683919 | 3.07E-19 |
| 0.37<=AIA<0.46 | 842 | 0.5472275 | 0.1008054 | 0.01739819 | 9.73E-09 |
| AIA<0.37 | 888 | 0.5369993 | 0.1161775 | 0.01815834 | 2.55E-10 |

**Supplementary Table 4:**
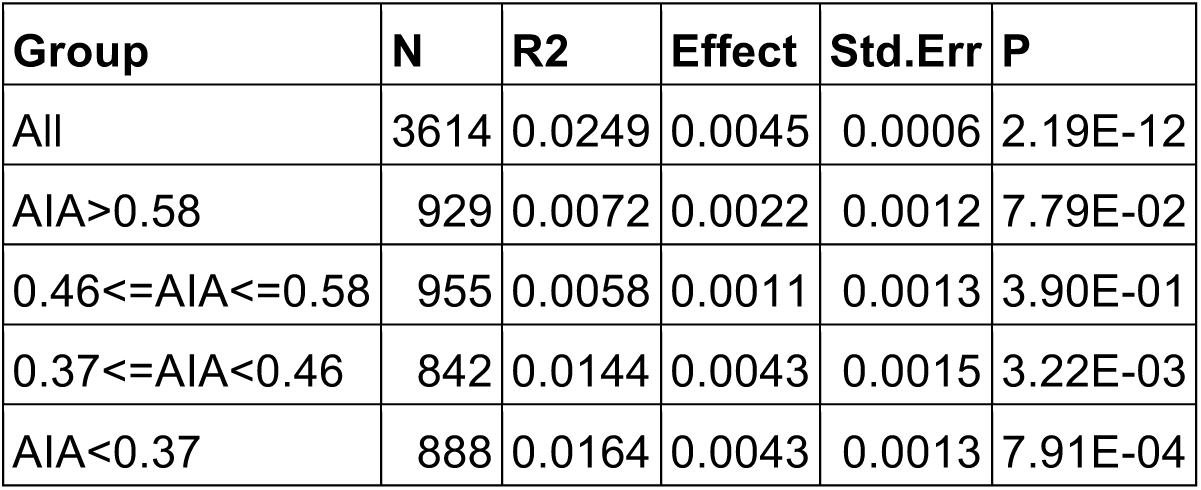
Predicted height vs. observed height. Predicted height (cm) as a function of observed height (cm) adjusting by sex, center, and sampling weight for 3614 Mexican Americans stratified by Amerindigenous ancestry (AIA).

| Group | N | R2 | Effect | Std.Err | P |
| --- | --- | --- | --- | --- | --- |
| All | 3614 | 0.0249 | 0.0045 | 0.0006 | 2.19E-12 |
| AIA>0.58 | 929 | 0.0072 | 0.0022 | 0.0012 | 7.79E-02 |
| 0.46<=AIA<=0.58 | 955 | 0.0058 | 0.0011 | 0.0013 | 3.90E-01 |
| 0.37<=AIA<0.46 | 842 | 0.0144 | 0.0043 | 0.0015 | 3.22E-03 |
| AIA<0.37 | 888 | 0.0164 | 0.0043 | 0.0013 | 7.91E-04 |

